# Reconstruction and analysis of the transmission network of African swine fever in People’s Republic of China, August 2018–September 2019

**DOI:** 10.1101/2020.07.12.199760

**Authors:** Andrei R. Akhmetzhanov, Sung-mok Jung, Hyojung Lee, Natalie Linton, Yichi Yang, Baoyin Yuan, Hiroshi Nishiura

## Abstract

Introduction of African swine fever (ASF) to China in mid-2018 and subsequent transboundary spread across Asia devastated regional swine production, affecting live pig and pork product-related markets worldwide. In order to explore the spatiotemporal spread of ASF in China, we reconstructed possible ASF transmission networks using nearest neighbour, exponential function, equal probability, and spatiotemporal case-distribution algorithms. From these networks we estimated the reproduction numbers, serial intervals, and transmission distances of the outbreak. The mean serial interval between paired units was around 29 days for all algorithms, while the mean transmission distance ranged from 332–456 kilometres. The reproduction numbers for each algorithm peaked during the first two weeks and steadily declined through the end of 2018 before hovering around the epidemic threshold value of one with sporadic increases during 2019. These results suggest that: 1) swine husbandry practices and production systems that lend themselves to long-range transmission drove ASF spread, and 2) outbreaks went undetected by the surveillance system. China and other affected countries have stepped up efforts to control ASF within their jurisdictions, and continued support for strict implementation of biosecurity standards and improvements to ASF surveillance are essential for halting transmission in China and further spread across Asia.

## Introduction

African swine fever (ASF) is a highly contagious and fatal disease of domestic pigs and wild boars (*Sus scrofa* spp.) that recently emerged in Asia. First described in Kenya in 1921 (Montgomery, 1921), ASF spread to countries in Western Europe during the mid-1950s but was painstakingly eliminated across the region with the exception of the island of Sardinia. ASF then made its way into Eastern Europe in 2007 via Georgia (Khomenko et al., 2013) before appearing in China in mid-2018, after which it infiltrated nearby Asian countries over the course of 2019 (FAO, 2019a). It is unclear how ASF entered China, but the circulating strain was genetically similar to the virulent strain of genotype II found in Georgia and Russia (Costard et al., 2013; Zhou et al., 2018). Although the ASF virus (ASFV) does not infect humans, control of its spread is of immense international concern as domestic pigs are a valuable source of food and other commodities. Pork products account for 37% of global meat consumption (FAO, 2019b), and products derived from pig parts are widely used in household and industrial applications, as well as for lifesaving medical treatments (Vilanova et al., 2019). Currently, there are no treatments or effective vaccines available to combat ASF.

Near 100% fatality of infected pigs has resulted in biosecurity measures that include the immediate culling of all pigs at sites where ASFV is detected, as well as blanket depopulation within 3-kilometre (km) epidemic zones around infected units (e.g., farms, backyards). Following such measures, Chinese authorities reported culling around 1.2 million pigs between August 2018 and June 2019 and it was reported that in August 2019 there was a −37.9% decrease in the number of live pigs in China compared to the same time the previous year (FAO, 2019c). Unfortunately, the multifactorial nature of ASF transmission precludes culling alone from halting disease spread.

For the most part, ASF transmission between swine occurs through direct contact or aerosol routes, but the virus can also be transmitted via arthropod vectors, fomites, or consumption of infected pork-related products—typically through the practice of feeding pigs raw swill, which is food scraps or food waste that contains or has come into contact with meat or meat products (EFSA et al., 2018). Indeed, early epidemiologic studies found that spread of ASFV in China was associated with biosecurity breaches such as raw swill feeding, improper disposal of dead animals, contaminated vehicles and workers, and transport of live pigs and their products across regions (MARA, 2019a; Wang et al., 2018). Armed with this knowledge, China banned the use of nonheated swill, disallowed swill feeding in provinces with ongoing outbreaks, set up inspection and disinfection stations to control farm traffic within 10-km buffer zones around the epidemic zones, limited the transportation of live pigs, closed and disinfected slaughterhouses in provinces with reported outbreaks, and closed live pig markets in affected and adjacent provinces (MARA, 2019b; Wang et al., 2018).

Identification of outbreak transmission events and development of effective control programs remain works in progress for affected countries (EFSA et al., 2018; Miller and Pepin, 2019). However, though infector-infectee relationships are largely unestablished, it is possible to identify likely infector-infectee pairs by reconstructing transmission networks using temporal and spatial data outbreak data reported by China to the World Organization for Animal Health (OIE). This data can also be used to quantify transmissibility of ASF spread by calculating the reproduction number *(R),* which represents the average number of secondary infections caused by a single, primary infection. When *R* is < 1, transmission is reduced and the number of infections decreases with each generation. Previous between-unit estimates of *R* during ASF outbreaks ranged from 2–3 between farms in Russia (Gulenkin et al., 2011) to 1.6–3.2 between pig herds in Uganda (Barongo et al., 2015).

In this study, we quantified factors of ASF transmission and surveillance over the course of the epidemic in China. First, by estimating the reporting delay—an indicator of how fast the outbreaks are being recognized by the authorities—and outbreak length—an indicator of how effectively each outbreak is managed. Next, we implemented four algorithms to reconstruct possible transmission networks and characterize the dynamics of ASF spread in China. This allowed us to examine changes in the ASF reproduction number over time as well as calculate the mean transmission distance and serial intervals (time between illness onset for pigs in infector-infectee paired units) of ASF in China.

## Results

As of 9 September 2019 there were 148 outbreaks of ASF in China. All provinces in mainland China experienced at least one outbreak (Figure 1; Figure 1—figure supplement 1). Outbreak units were classified as farms, backyard farms, villages, or slaughterhouses. The size of pig population supported varied between and within unit types (Figure 1—figure supplement 2). We found no significant correlation between outbreak start date and affected unit size (Pearson’s *r* = −0.044, *N* = 147, *P* = 0.589) or type (ANOVA, *F*(3,144) = 1.96, *P* = 0.123), nor between the fraction of infected pigs and live pig density by province (Pearson’s *r* = −0.15, *N* = 147, *P* = 0.406).

**Figure 1.**
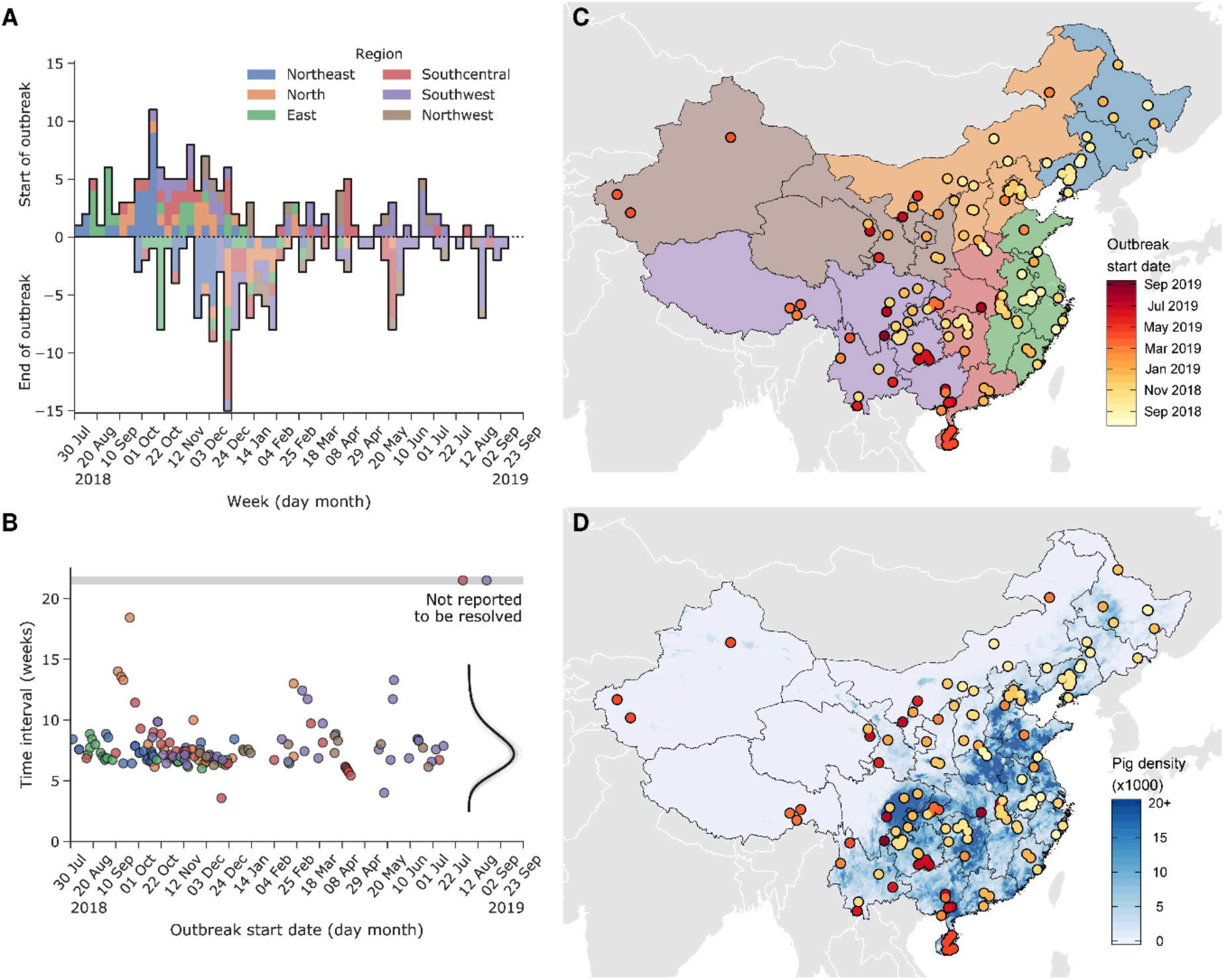
Characteristics of African swine fever (ASF) infected farms in China from July 2018–May 2019. (A) Weekly number of reported outbreaks by outbreak start and end dates for the six regions in China. (B) Time interval between the outbreak start and end dates by date of outbreak start. The point colours represent the region in which each infected unit is located, consistent with colours in (A). Points within the horizontal grey bar are unresolved cases. Inset in (B): The right-hand vertical line with grey shading indicates the distribution of the time interval and 95% confidence intervals, respectively. The scale is not shown, but the area under the curve is equal to one. (C) Geographical distribution and outbreak start date of ASF infected farms. Point colours indicate the start date of outbreak in each infected unit. (D) Pig density and geographical location of ASF infected unit. Point colours indicate the start date of outbreak and blue shade presents the density of lived pigs in China, as reported by the Food and Agriculture Organization (FAO).

Reporting delay for the outbreaks had a mean of 7.6 weeks (95% credible interval [CI]: 7.3–7.8) and a standard deviation (SD) of 1.6 weeks (95% CI: 1.5–1.8) (Figure 1B; Figure 1—table supplement 1). The distribution of the reporting delay displayed a heavy tail and bimodality (Figure 1—figure supplement 3). We fit the reporting delay to a combination of lognormal (for shorter reporting delays) and gamma (for longer reporting delays) distributions, as this combination yielded the minimal median Watanabe-Akaike information criterion (WAIC) value (Figure 1—table supplement 1). The mean reporting delay assuming a shorter reporting delay (lognormal distribution) was 5.4 days (95% CI: 4.4–6.5) with an SD of 3.8 days (95% CI: 2.3–5.5), and the mean reporting delay assuming a longer reporting delay (gamma distribution) was 21.6 days (95% CI: 6.8–34.2) with an SD of 10.7 days (95% CI: 0–21.0). The mean reporting delay using the combined lognormal and gamma distributions was 6.3 days (95% CI: 5.5–7.2) with a SD of 5.3 days (95% CI: 4.1–6.7). There was no clear correlation found between reporting delay and unit type (ANOVA, F(3,131) = 1.25, *P =* 0.293) or unit size (Pearson’s *r* = −0.04, *N* = 134, *P* = 0.655) (Figure 1— figure supplement 2AB).

Figure 2 shows the reconstructed transmission networks of ASF spread among infected units using nearest neighbour, exponential function, and equal probability algorithms, which rely on knowledge of the geographic location of reported outbreaks—see Materials and Methods section for details. We required that the time between paired outbreak start dates was no shorter than the ASF incubation period. The nearest neighbour algorithm yielded an estimated mean transmission distance of 332 km (95% CI: 213–1548 km) with a mean serial interval of 29.0 days (95% CI: 6–62 days) between paired transmission events. The exponential function algorithm yielded a mean transmission distance of 456 km (95% CI: 22–1550 km) and a mean serial interval of 29.3 days (95% CI: 6–64 days) with the distance kernel set at 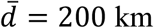. Due to the time constraint we imposed, we were unable to link an average of 8 outbreaks (95% CI: 5–12) to any potential infector for these two methods (Figure 2—figure supplement 1AB).

**Figure 2.**
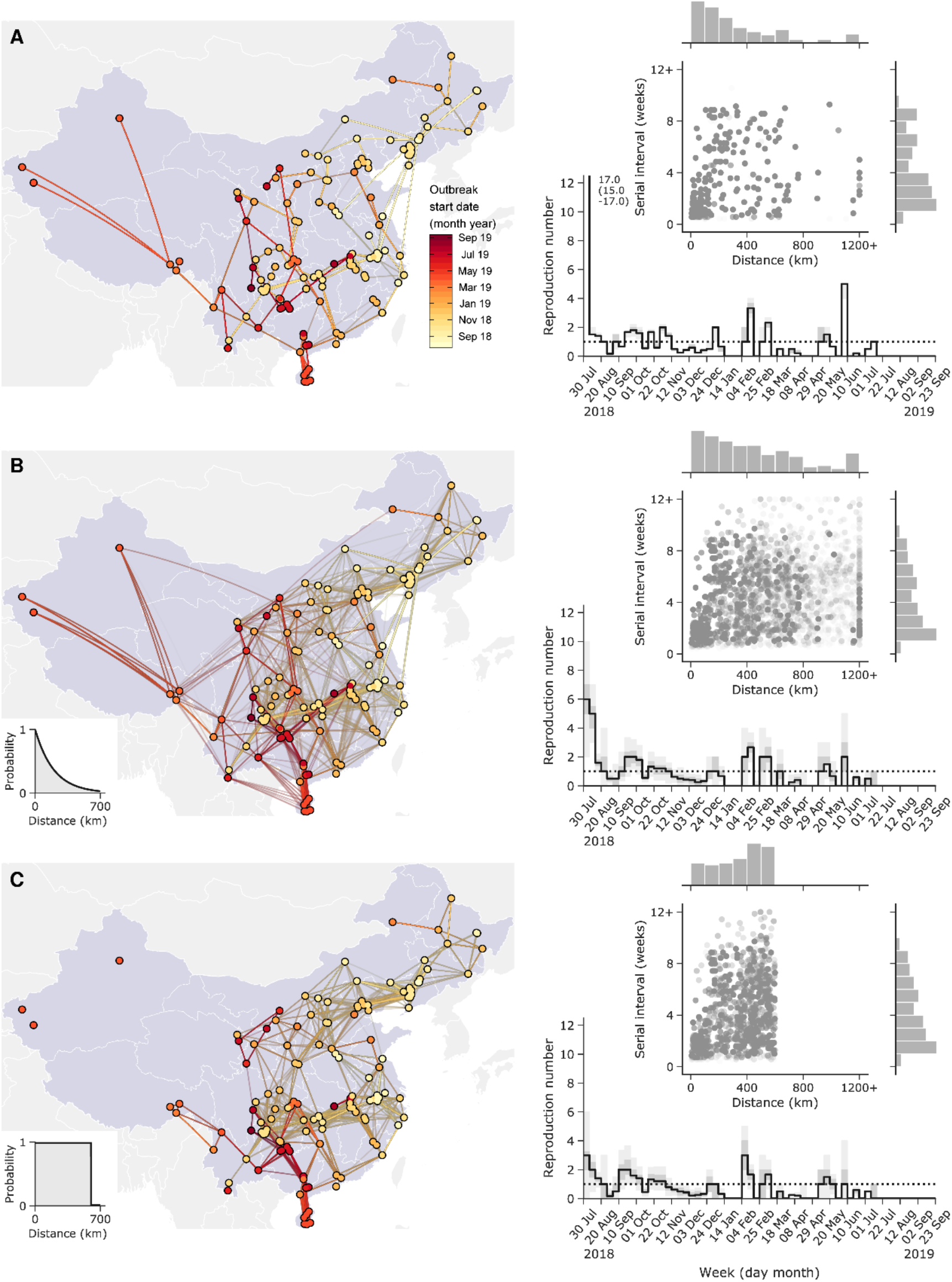
Reconstructed transmission networks of African swine fever (ASF) outbreak from July 2018–September 2019 in China and estimates of reproduction number and serial interval from reconstructed networks. Three transmission networks are reconstructed by using (A) nearest neighbour, (B) exponential function and (C) equal probability algorithms, analysing only outbreaks reported to the World Organization for Animal Health (OIE). The dot and line colours in the map represent the start date of the outbreak in each infected unit. Correlations between the serial interval and transmission distance are shown in upper right side of each figure. The points indicate each ASF infected farm and bars represent distribution of estimated distance and serial intervals, using the reconstructed transmission networks, respectively. The lines and shades in each of the figures on the right show the estimated reproduction number and its 95% confidence intervals by calendar week. The epidemic threshold (*R* = 1) is represented with a dashed line.

The equal probability algorithm yielded a mean transmission distance of 344 km (95% CI: 23–595 km) and mean serial interval of 29.5 days (95% CI: 6–64 days) with the effective distance set at 600 km, which is the 95th percentile of the previous kernel function. An average of 30 outbreaks (95% CI: 28–34; Figure 2—figure supplement 1CD) could not be linked to any potential infector due to the constraint on long-range transmission imposed by setting the effective transmission distance at 600 km. We performed sensitivity analyses for effective transmission distance used in the exponential function and equal probability algorithms, but did not find that either algorithm was sensitive to the value of the effective transmission distance (Figure 2—figure supplements 2–3).

Our use of the spatiotemporal case-distribution algorithm developed by Salje and colleagues (Salje et al., 2016) resulted in a mean transmission distance of 483 km (95% CI: 449–503 km). A variation in the mean (range 1–14 weeks) and SD (range 1–7 weeks) of the generation time distribution led to the approximate range of the mean transmission distance between 200 and 500 km (Figure 3). The mean transmission distance tended to increase if either the mean or SD of the generation time were increasing. Additional numerical simulations of a virtual outbreak of ASF (Appendix II) revealed that removal of a fraction of reported cases with both temporal and spatial information led to a substantial overestimation of the mean transmission distance, as shown in Appendix Figure 2.

**Figure 3.**
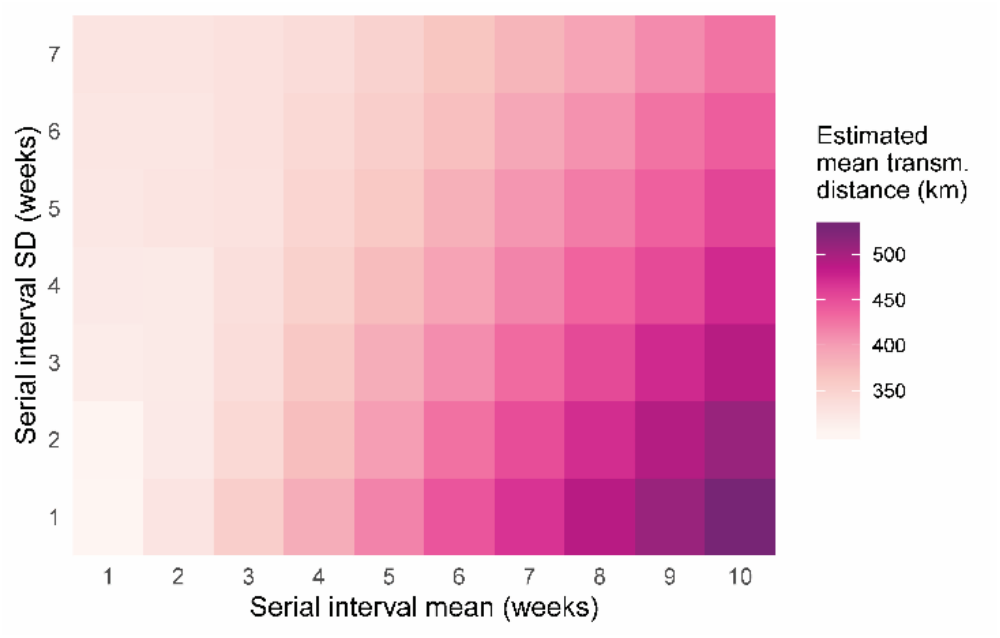
Mean transmission distance based on varying mean and standard deviation (SD) values of the serial interval distribution. Estimation relies on a generalized Wallinga-Teunis method developed by Salje and colleagues (Salje et al., 2016). Both the spatial transmission kernel and serial interval are assumed to follow a normal distribution, with 1000 simulations of the transmission networks used for each particular value of the mean and SD. For additional details see the Materials and Methods section.

## Discussion

ASFV remains in circulation in China more than one year since it was first detected. To our knowledge, this is the first study to use mathematical modelling to quantify ASF transmission within China. We found that the median serial interval was approximately 29 days and the mean distance between suspected transmissions ranged from 332–456 km, reflecting the wide temporospatial spread of the epidemic in China. These results suggest that: 1) swine husbandry practices and production systems that lend themselves to long-range transmission drove ASF spread, and 2) outbreaks went undetected by the surveillance system.

The initial reduction of the ASF reproduction number below the threshold value of one for our three geographically-based algorithms coincided with official reports of successful outbreak control measures in November 2018. However, persistent transmission led to sporadic increases in the reproduction number later in 2019. The swiftest decline in the reproduction number occurred following the nearest neighbour algorithm, while the exponential and equal probability models exhibited smoother, transient dynamics. Having now been detected in many countries in Asia, ASF has become a critical international biosecurity concern and our estimates of the weekly reproduction number, serial interval, and transmission distance of ASF in China can help inform intervention and management strategies.

Outbreak underascertainment is the main limitation of our study. It is plausible that ASF surveillance capacity in China was not sensitive enough to detect all infections— particularly in smaller units with lower biosecurity—leading to underascertainment of outbreaks and consequently underestimation of the reproduction number with the methods we used. We suspect that the large estimates of the mean transmission distance could be a consequence of underascertainment of infected units. The estimates of the mean transmission distance using the spatiotemporal case-distribution method were also larger than anticipated. This was likely due to underascertainment of infected units or intentional culling of susceptible pigs in within the epidemic zones surrounding the reported units. Both factors would reduce the number of susceptible pigs within a shorter transmission distance, therefore making transmission of ASF to new regions possible only through long-range transmissions and prolongated generation intervals. Although we investigated the possible impact of outbreak underascertainment on transmission distance using simulation data, there are still some methodological uncertainties when both temporal and spatial information is missing. If the true degree of underascertainment can be estimated (e.g., using genetic data or environmental sampling) the ASF reproduction number could be more accurately assessed.

In our study, we also did not consider importation of ASF virus from outside of China, and only modelled within-country transmission. This could perhaps be accomplished using other data, but is beyond the scope of this study. We also did not fully consider the impact of the lunar New Year—one of the biggest holidays in China—in our calculations of the reporting delays and outbreak lengths. However, we could not find any clear correlation between the submission delay (i.e., time interval between notification date of outbreak in Chinese government and report submission date to OIE) and official dates of the holiday.

Control of ASF in China, the world’s leading producer and consumer of pork (USDA, 2019), is of critical importance to countries that import Chinese pig products as the virus may remain viable in blood and tissues for long periods of time (Plowright and Parker, 1967; Sindryakova et al., 2016). Although vaccines for ASF are under development (Barasona et al., 2019), biosecurity-based control measures remain key to preventing the transboundary spread of ASF, and better understanding of the transmission distances and transmissibility of ASF in China can help inform management strategies to prevent further spread. The disease poses an ongoing threat to livelihoods and national swine-related gross domestic product, as well as to the development of important medical, industrial, and household products.

Although China has endeavoured to contain the spread of ASF, the economic implications of the outbreak have begun to show. Chinese import of pork from the United States is well above 2018 levels (Sánchez et al., 2019) and the lifesaving drug heparin, mainly produced by Chinese companies from porcine mucosa, may be in short supply

(Vilanova et al., 2019). The various transmission pathways for ASF and inherent biosecurity risks and economic devastation to small-scale farms remain a persistent concern for China and neighbouring countries. After examining four algorithms to assess transmission pathways, we found a large mean transmission distance and lengthy serial intervals, which are likely due to underascertainment of cases and a prevalence of long-range transmission events. Our results indicate that continued monitoring of transmission, improvements to surveillance across large distances and lengthy time periods, and increased biosecurity measures are critical to eradication of ASF in China. As well, it is important for China and other countries to consider all possible modes of transmission when developing biosecurity protocols, as wild boar reservoirs and arthropod vectors may further complicate transmission networks. Restructuring of biosecurity protocols are vital to containment of the threat at national and international levels.

## Materials and Methods

### Epidemiological data

The dataset of all ASFV infected units was aggregated from publicly available immediate notifications and follow-up reports submitted by the Ministry of Agriculture and Rural Affairs of the People’s Republic of China (MARA; http://www.moa.gov.cn) to OIE beginning in August 2018 and published in the World Animal Health Information System (WAHIS) https://www.oie.int/wahis_2/public/wahid.php/Wahidhome/Home). An ASF outbreak was defined as at least one pig infected with ASFV within a single unit—either a farm, backyard, slaughterhouse, or village. In China, detection of ASF triggers the establishment of an epidemic zone with a radius of 3 km around the infected unit within which all pigs are culled and entrance of live pigs is restricted by blockade. A 10-km buffer zone is also established, in which inspection and disinfection stations are set up to control traffic to and from the infected unit. Infected slaughterhouses were shut down and decontaminated, and MARA now tasks slaughterhouses with conducting self-inspection using laboratory tests (MARA, 2019c). Live pig trading markets in affected areas were closed. Outbreaks related to infected units were declared over when there were no new infections within the epidemic zone for 6 weeks (MARA, 2019d). End of outbreak information was not retrospectively updated in the WAHIS reports, so we supplemented the data with dates retrieved from official announcements by MARA. If the dates obtained from WAHIS and MARA for the same outbreak differed, we used the earlier date. To assess spatial correlation between the density of live pigs and the location of infected units, we used values of the global livestock population predicted by the Gridded Livestock of the World (GLW v3) system of the FAO of the United Nations (http://www.fao.org/livestock-systems/en/).

A total of 155 outbreaks were reported between 4 August 2018 and 9 September 2019, of which 152 were outbreaks among domestic pigs, two were outbreaks on wild boar farms in Heilongjiang and Inner Mongolia, and one was a report of a sole infected wild boar in Jilin Province. The wild boar was excluded from our analyses. We also disregarded six outbreaks reported in March, June, and August 2019 linked to interception of transport vehicles carrying infected pigs. In the first two reports, transport trailers carrying 150 pigs (9 dead) and 32 pigs (1 dead) were intercepted at highway checkpoints for animal health supervision in Sichuan and in Guizhou provinces and the origin of these pigs was uncertain (Guang’an Municipal People’s Government, 2019). Swine in the other four outbreaks in late August 2018 similarly had unknown origin. Exclusion of these reports resulted a total of 148 outbreaks included for analysis.

### Reporting delay

We defined reporting delay as the time between the date of outbreak start and the date of outbreak notification to OIE. Among the 148 outbreaks, only 135 included both the start and notification dates. We right-censored the records with unspecified report date and assumed that the reporting date could be any time between one day after the start of the outbreak and the date of release of the report by OIE. We estimated the distribution of the reporting delay using a discretized probability distribution derived from the cumulative distribution functions *H*(*t*; ***α***) of single gamma, lognormal, and Weibull distributions, and their mixtures. The probability an infected unit was reported at time *t* after infection is detected is modelled by: *h_t**α**_* = *H*(*t* + 1; ***α***) — *H*(*t; **α***). The parameter ***α*** represents a vector of the means and SDs of the distributions. In case of a mixture, the function *h_t **α**_* is decomposed into two discretized probability distributions, as: *h*_t,**α**_ = *αh*_*t***α**_1__ + (1 – *ρ*)*h*_*t****α***_2__, where *ρ* is the relative weight of the first distribution over the second (0 < *ρ* < 1); ***α*** = {***α***_1_, ***α***_2_, ρ}, **α**_*i*_ = {*μ_i_*, *σ_i_*} consists of the mean (*μ_i_*) and SD (*σ_i_*) of the probability mass function *h_t,α_i__* (*i* = 1, 2).

### Reconstruction of probable transmission networks

We employed three transmission kernels *f* following nearest neighbour, exponential, and equal probability algorithms (Kraemer et al., 2017; Riley et al., 2015; Viboud et al., 2006) to reconstruct probable transmission networks of ASF in China. For each algorithm, *f* is dependent solely on the distance between paired units. For the nearest neighbour kernel, the unit closest to the infectee was selected as the infector. In contrast, the exponential and equal probability kernels have an exact functional form, such that the exponential kernel 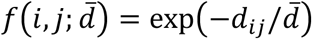 where 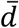 is the scale of the effective transmission distance, and the equal probability kernel 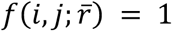 if 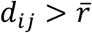 and is 0 otherwise. The two kernels will produce an identical transmission network if 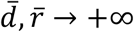. For the equal probability algorithm we set the probability of transmission as equally likely within a given radius 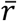 and zero otherwise. In our simulations we set 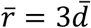 which is also equal to the 95th percentile of the exponential kernel function. All algorithms restricted unit linkage to infector and infectee pairings where the estimated serial interval was longer than the ASF incubation period based on a gamma distribution with a mean of 6.3 days and SD of 1.3 days (Olesen et al., 2017).

Next, we assigned weights *w_ij_* to each potential infector *i* and infectee *j* pairing. The value for each weight was set to the respective value of the kernel function *f*(*i,j*|°) unless the following two conditions were violated: (i) the outbreak began at the infectee location later than it began at the infector location plus the estimated incubation period for ASF; and (ii) the outbreak began earlier than it ended at the infector location minus the incubation period. Otherwise, the weight *w_ij_* was set to zero. The incubation period distribution was given as a gamma distribution with a mean of 6.3 days and SD of 1.3 days (Olesen et al., 2017). Imputed outbreak end dates (*n* = 2, as in Figure 1B) were drawn from a gamma distribution of all known outbreak start to end time intervals. Implementation of these two uncertainties resulted in a probabilistic non-uniqueness of the transmission network even when linkage of an infector and an infectee was established using the nearest neighbour algorithm.

The transmission probabilities *p_ij_* for each infectee were set by normalizing the weights for all potential infectors as: 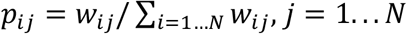. The infectee is linked to a single infector by sampling from the categorical distribution:

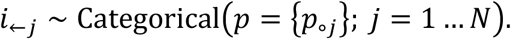

When *p_ij_* = 0, linkage of infectee to infector does not occur (e.g., the index outbreak).

Lastly, we estimated the mean transmission distance based on the methods developed by Salje and colleagues (Salje et al., 2016), which extends the temporal framework developed by Wallinga and Teunis (Wallinga and Teunis, 2004) to include a spatial dimension. The temporal dimension is governed by the generation time distribution, and the spatial dimension is characterized by a transmission kernel distribution which describes a probability of observing two cases at given distance. We used a probability generation function *g*(*t; θ_g_*), where the parameter set *θ_g_* = {*μ_g_, σ_g_*} consists of the mean *μ_g_* and standard deviation *σ_g_* to describe the generation time distribution. The distribution was discretized by day following the form of: 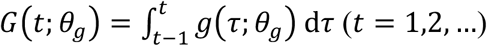. Next, we defined a Wallinga-Teunis matrix 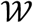 where rows represent infectees *i* and columns represent potential infectors *j*. Each element of the matrix is a probability of transmission between *i* and *j* of

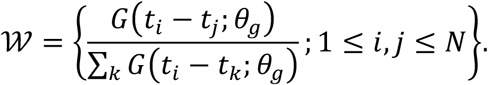

The infectee-infector pairings (*i,j*) were identified using sampling from a categorical distribution respective to the rows of the Wallinga-Teunis matrix: 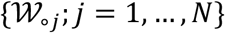. If all elements in a row of the matrix 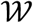 appeared to be zero, the case was left unpaired (e.g., the index outbreak).

We then used categorical sampling to construct the transmission network 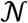. For any pair of nodes *i* and *j* we determined a set of network paths connecting them and recorded the number of links *ζ_ij_* in each path. This procedure was done repeatedly over a large set of sampled transmission networks to form the set of counts **Z**_*ij*_ = {*ζ_ij_*}. We then defined *π*(*ζ; i,j*) as the frequency of observing the number of links *ζ* between *i* and *j* in a set **Z**_*ij*_ and applied the formula obtained by Salje and colleagues to estimate the mean transmission distance by taking it as equal to the SD of the transmission kernel:

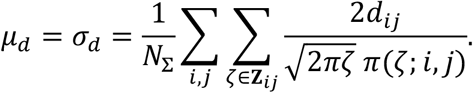

Here, *d_ij_* is the geographic distance between *i* and *j* and *N*_Σ_ is a total number of observed pairs (*i,j*).

---

**Figure 1—figure supplement 1.**
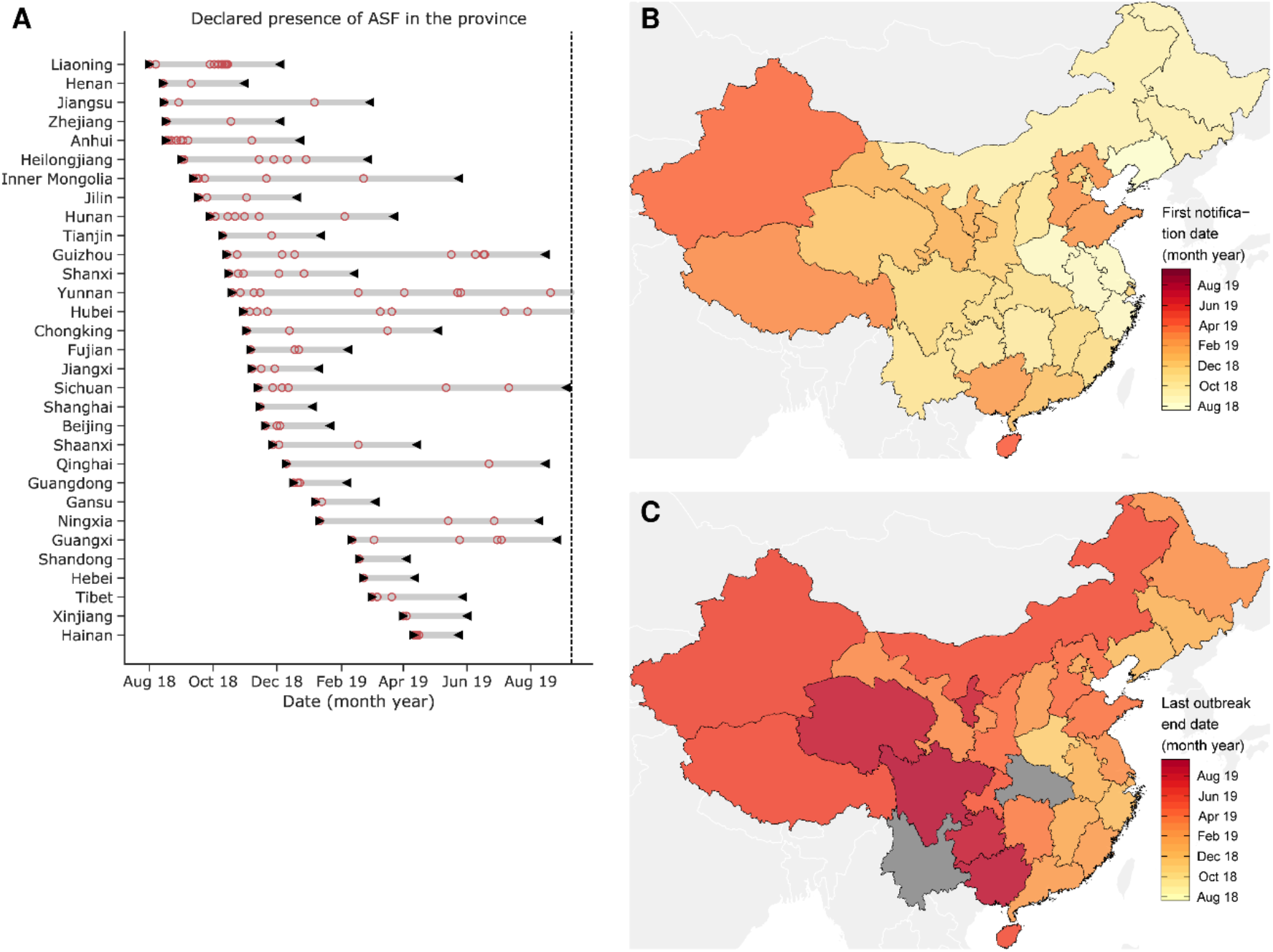
First appearance and last outbreak by province of China. (A) shows start date of the first outbreak and end date of the last outbreak-if declared-for each province (black triangles). The red circles indicate the start dates of the other outbreaks. (B) and (C) are heatmaps of first outbreak start date (B) and last outbreak end date (C) by province. The grey areas indicate provinces with ongoing outbreaks as of 9 September 2019.

**Figure 1—figure supplement 2.**
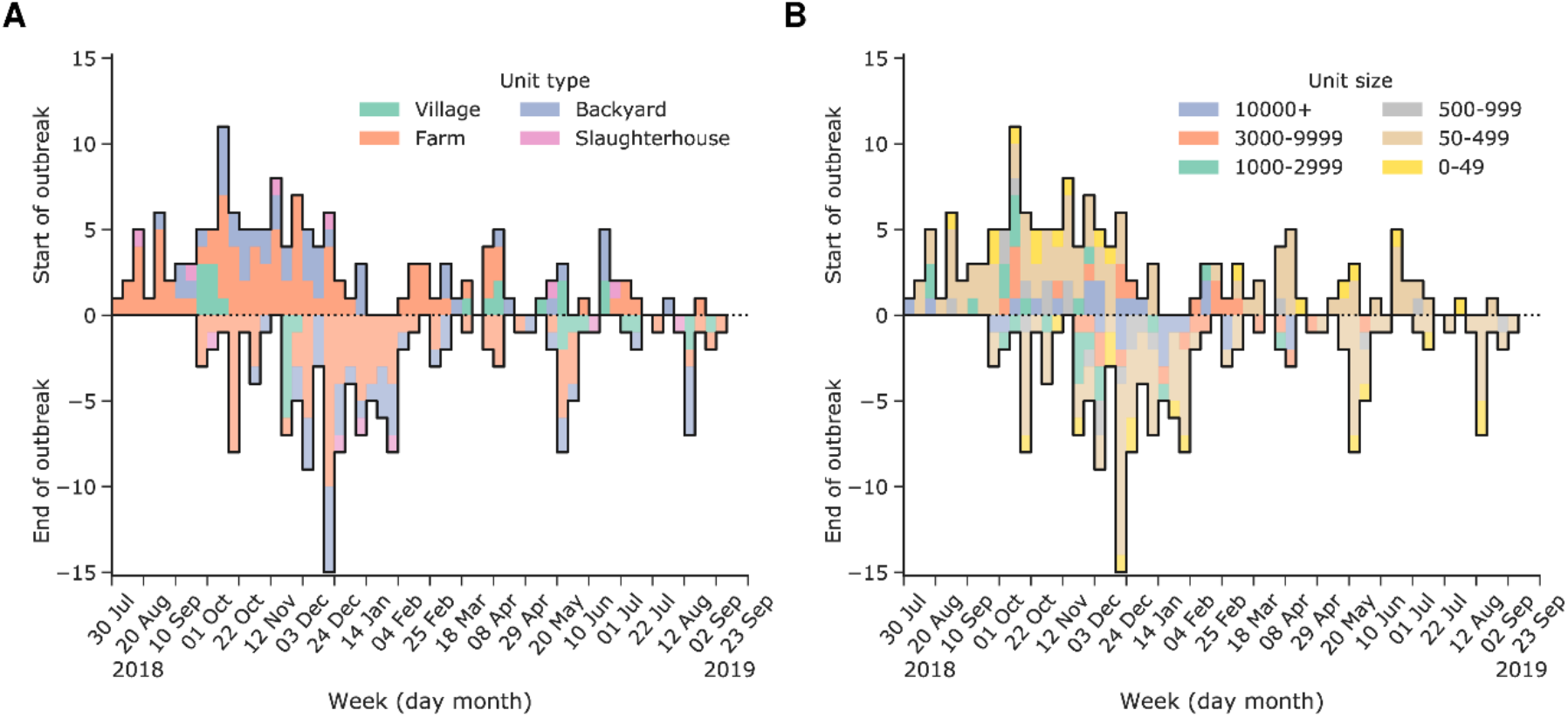
Weekly number of reported and terminated outbreaks of African swine fever (ASF) factorized by type (A) or size of the unit (B). Epidemic curves by (A) type of unit and (B) size of the unit are shown. The coloured bars indicate the different types and sizes of infected units in China from July 2018–September 2019.

**Figure 1—figure supplement 3.**
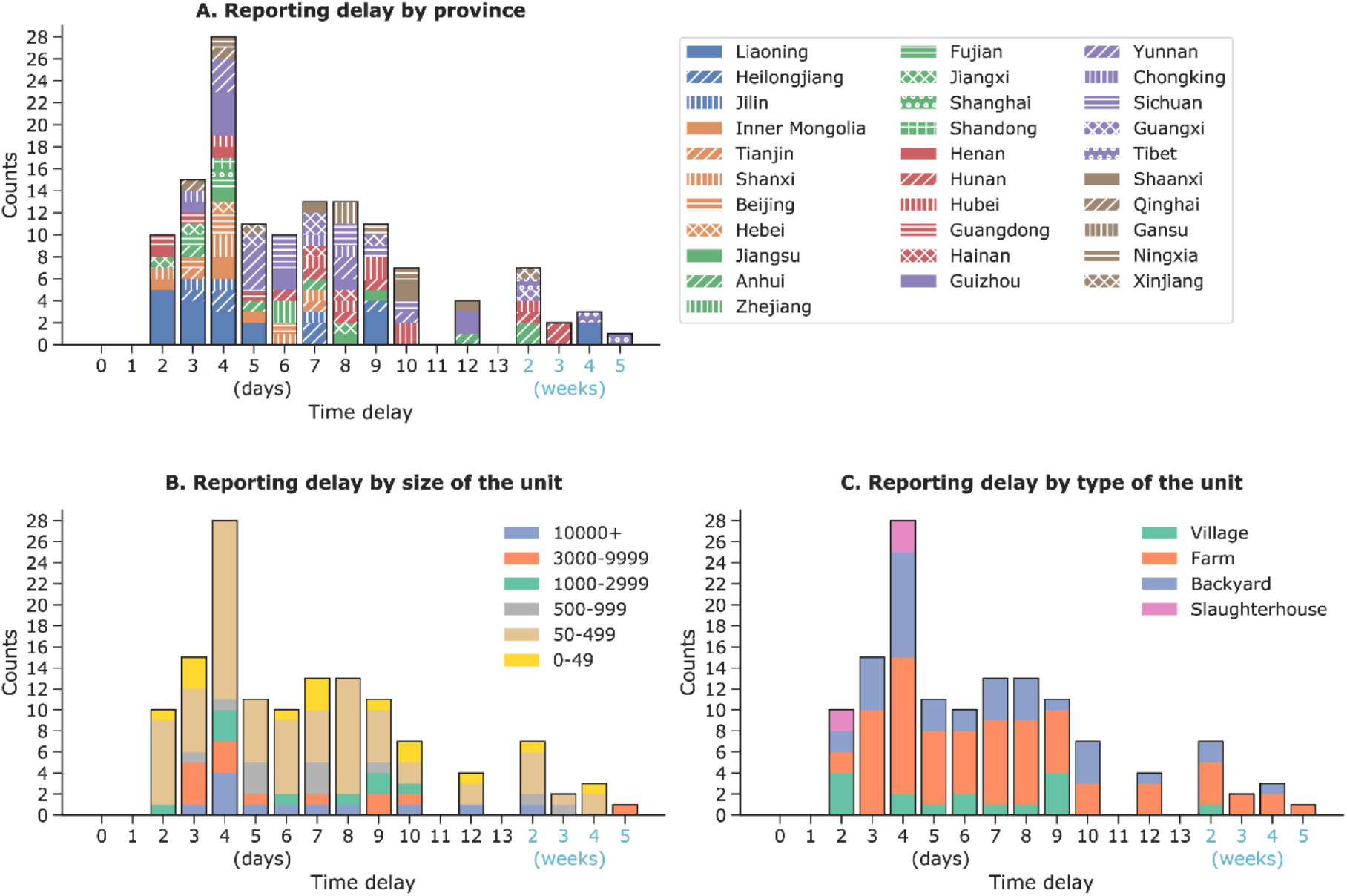
Reporting delay by characteristic of African swine fever (ASF) infected unit. Distribution of the reporting delay by (A) province, (B) size of the unit and (C) type of the unit are shown. Reporting delay is calculated using outbreak start and report dates, as released by the World Organization for Animal Health (OIE). The bar colours represent the various characteristics of the ASF infected units.

**Figure 1—table supplement 1.**
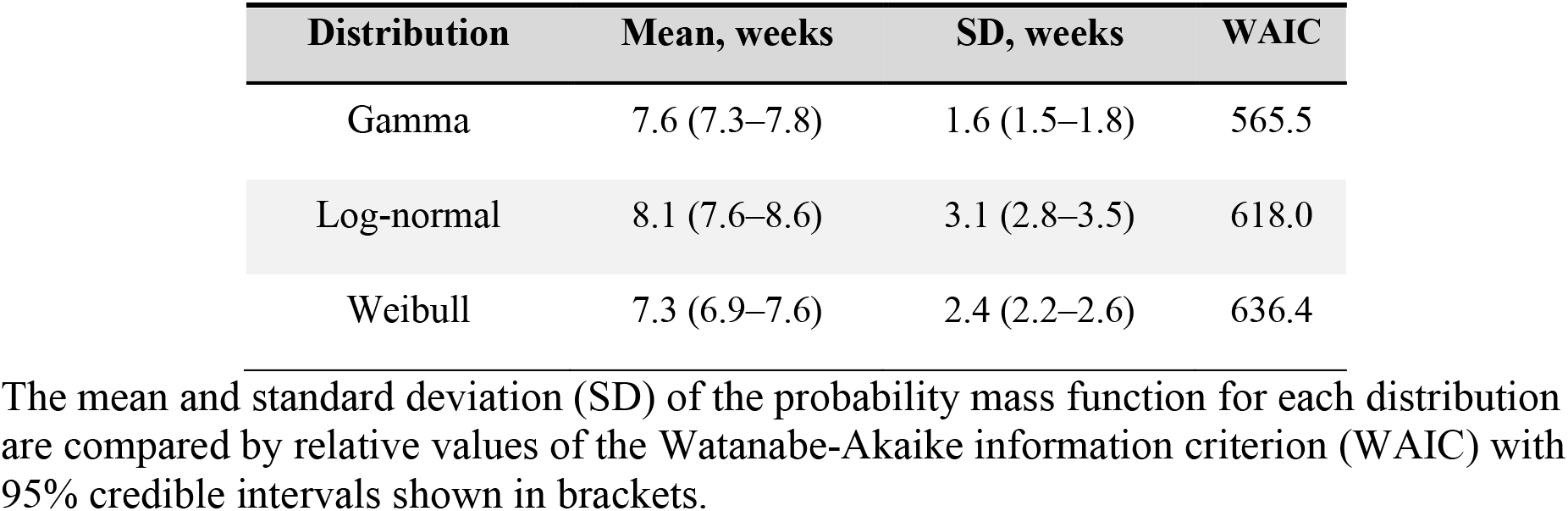
Fit of the time period between the start and the end of the outbreak to different distributions

| Distribution | Mean, weeks | SD, weeks | WAIC |
| --- | --- | --- | --- |
| Gamma | 7.6 (7.3–7.8) | 1.6 (1.5–1.8) | 565.5 |
| Log-normal | 8.1 (7.6–8.6) | 3.1 (2.8–3.5) | 618.0 |
| Weibull | 7.3 (6.9–7.6) | 2.4 (2.2–2.6) | 636.4 |

The mean and standard deviation (SD) of the probability mass function for each distribution are compared by relative values of the Watanabe-Akaike information criterion (WAIC) with 95% credible intervals shown in brackets.

**Figure 1—table supplement 2.** Fit of the reporting delay to different distributions

| Distribution | $\mu_1$, days | $\sigma_1$, days | $\mu_2$, days | $\sigma_2$, days | $\rho$ | WAIC |
| --- | --- | --- | --- | --- | --- | --- |
| Mixture of log-normal (1) and gamma (2) | 5.5<br>(4.4–6.5) | 3.9<br>(2.4–5.5) | 22.1<br>(8.4–35.0) | 10.9<br>(0.1–21.7) | 0.91<br>(0.77–1.00) | 793.8 |
| Single log-normal | 6.3<br>(5.5–7.2) | 5.3<br>(4.0–6.6) | — | — | — | 797.6 |
| Mixture of gamma (1) and log-normal (2) | 4.4<br>(2.3–6.1) | 3.0<br>(0.1–6.3) | 11.1<br>(5.5–23.7) | 9.8<br>(4.0–20.7) | 0.49<br>(0.00–0.91) | 799.1 |
| Mixture of two gammas | 4.9<br>(4.2–5.7) | 2.8<br>(2.0–3.5) | 18.7<br>(9.0–29.6) | 10.8<br>(5.3–18.3) | 0.85<br>(0.68–0.97) | 799.4 |

Parameters $\mu_i$ and $\sigma_i$ ( $i = 1, 2$ ) represent the mean and standard deviation of the probability mass function for each distribution of the reporting delay by distribution type, $\rho$ is the weight attributed to each configuration by the relative values of the Watanabe-Akaike information criterion (WAIC), and 95% credible intervals are shown in brackets. Although all possible combinations of single gamma, log-normal, Weibull distributions and their mixtures were considered, only the top four configurations with minimal WAIC values are shown here.

**Figure 2—figure supplement 1.**
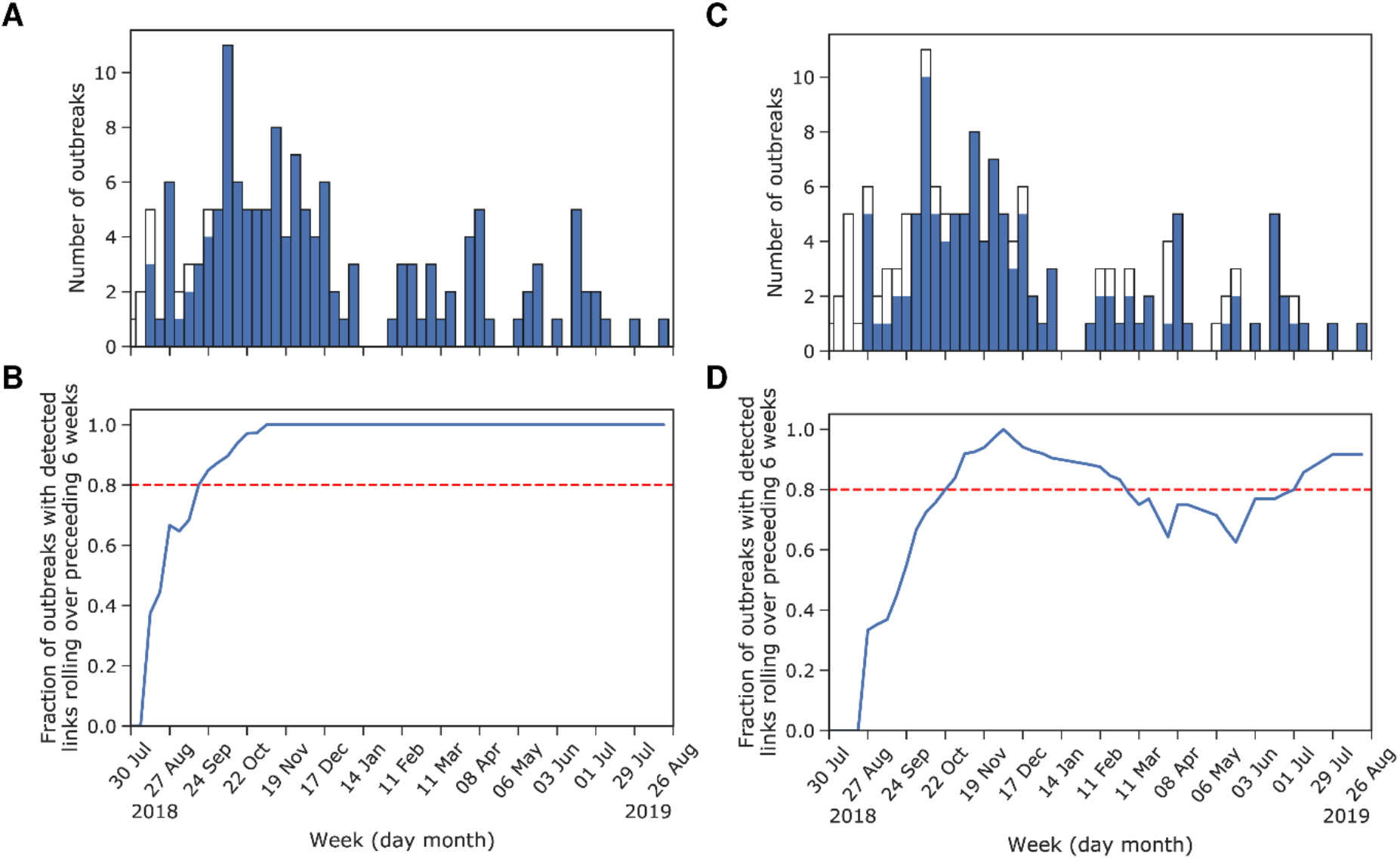
Outbreak linkage by nearest neighbour or exponential function algorithms (A, B) or by equal probability algorithm (C, D). (A) and (C) show the number of outbreaks with detected links (solid bars) vs total number of outbreaks by week of outbreak start. (B) and (D) show the fraction of outbreaks with detected links over the preceding six weeks by week of outbreak start. The dashed horizontal red line indicates a 80% fraction of detected links.

**Figure 2—figure supplement 2.**
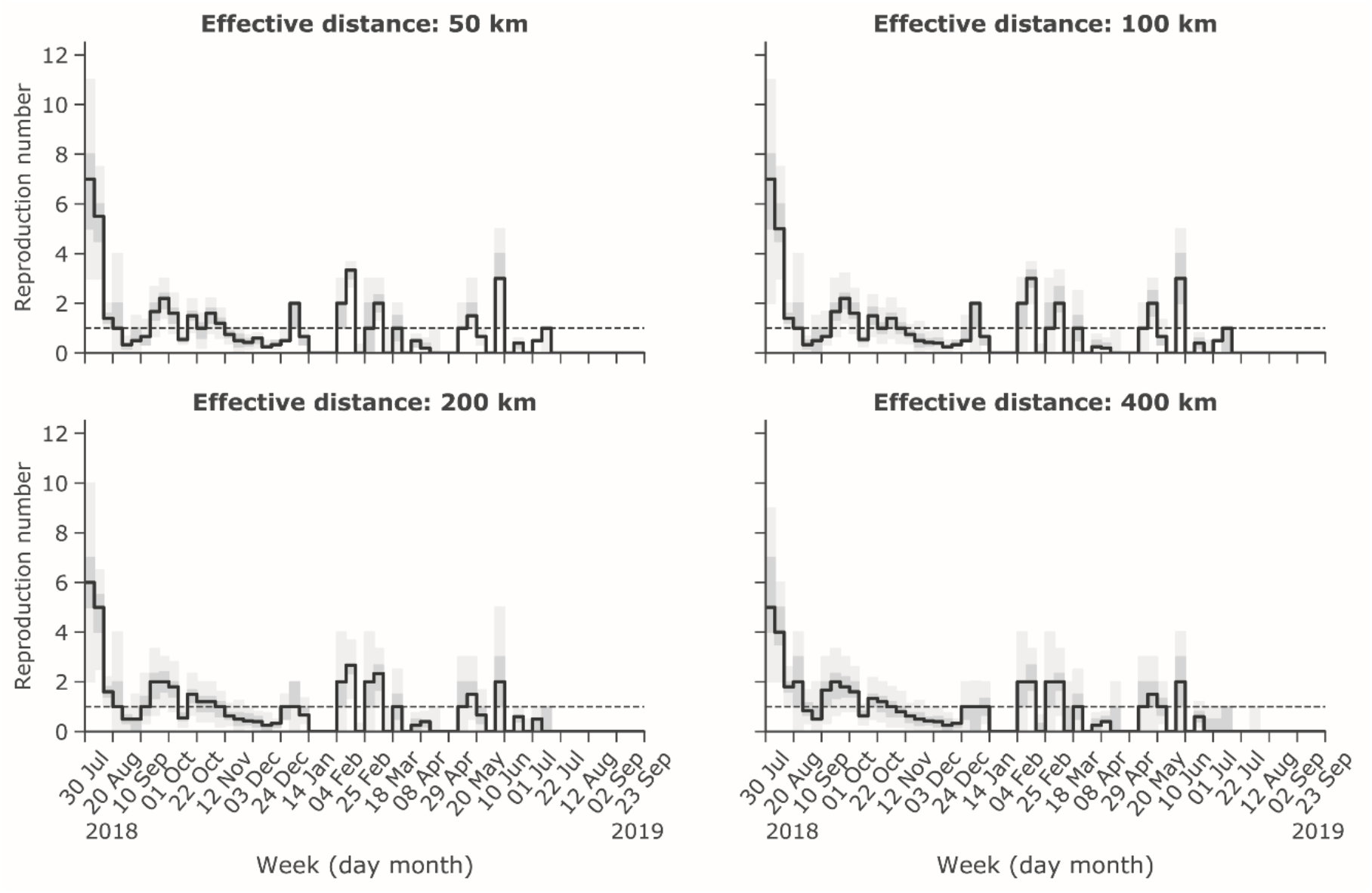
Estimated weekly reproduction number via exponential function algorithm using four different effective transmission distances 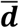. The lines and grey shading represent the reproduction number by week including 95% credible intervals. The threshold number of one is represented with a dashed line.

**Figure 2—figure supplement 3.**
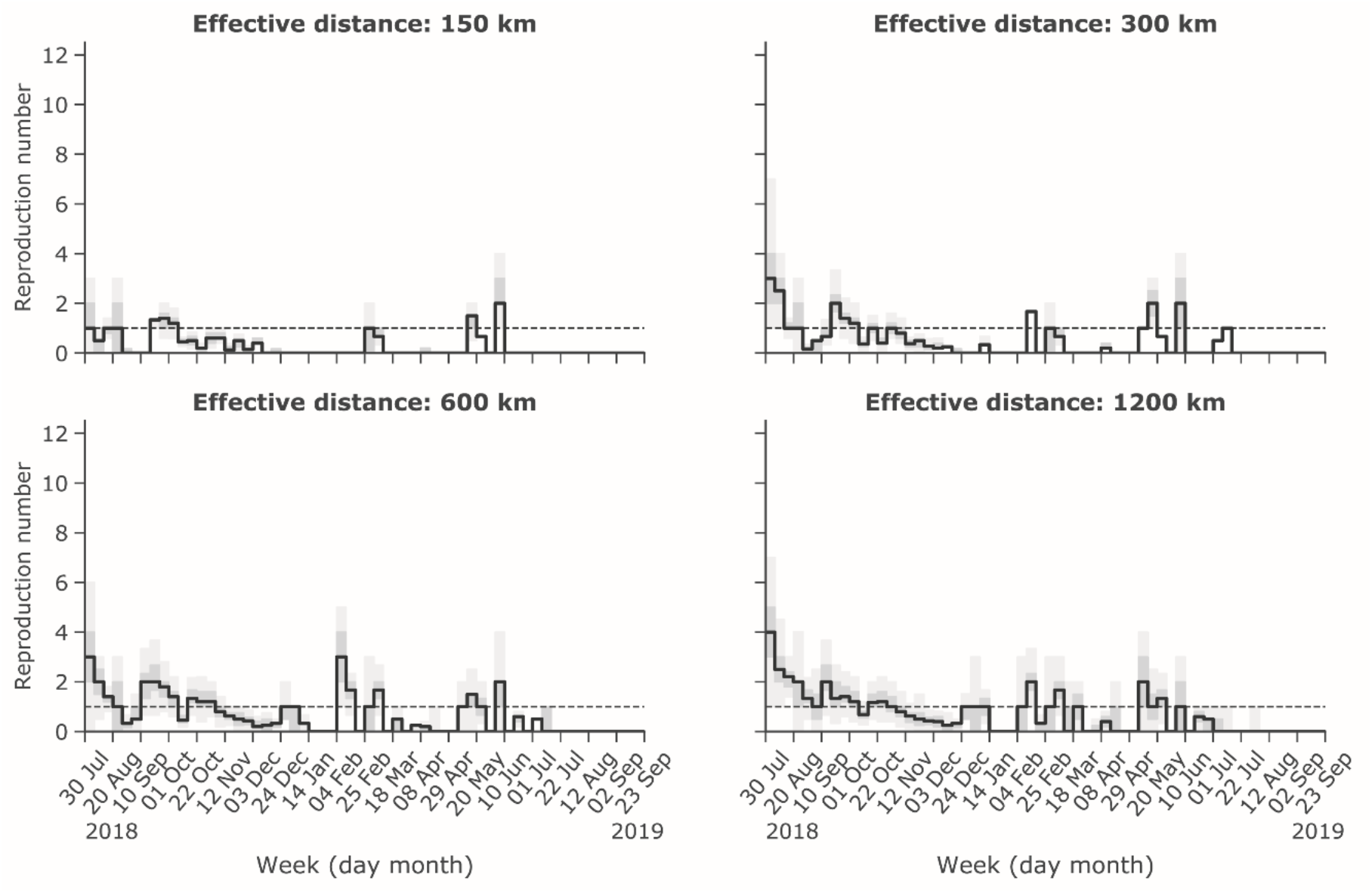
Estimated weekly reproduction number via equal probability algorithm, using four different effective transmission radii 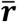. The lines and grey shading represent the reproduction number by week including 95% credible intervals. The threshold number of one is represented with a dashed line.

## Supplementary materials

### Appendix I. Inference of the generation time from historical records

Estimation of the mean transmission distance using the method of Salje and colleagues required knowledge of the generation time distribution of ASF between-farm transmission. As detailed investigation reports for the outbreaks in China were unavailable (USDA Foreign Agricultural Service, 2019) we utilized an investigation report from an ASF outbreak in the Odessa region of the former United Soviet Socialist Republics (USSR) in 1977 (Korennoy et al., 2017; SSI VNIIVViM, 2014) to obtain a between-farm generation time distribution.

Prior to re-introduction of ASFV into post-USSR territory (Georgia) in 2007, the first and only epidemic of ASF in the USSR was in the Odessa region of the Ukrainian Soviet Socialist Republic, a part of contemporary Ukraine, in 1977 (Appendix Figure 1A). The epidemic began around the end of February and continued until July. Twenty units were reported to be infected and a total of 360,500 pigs in epidemic zones and high-risk regions were culled, leading to a loss of 30.5% of pig herds in the region. The affected units ranged from small private backyards with only a few pigs to large collective farms with 1,509– 13,865 pigs. The main driver of spread was insufficiently preheated swill feed (Appendix Figure 1B).

Data were aggregated into weekly counts of infected units {*n_w_; w* = 1,2, … } with the first week set to the date of start of the index outbreak. The force of infection *λ_w_* was modelled using a sequential generation process referred to as a generation-dependent model (Akhmetzhanov et al., 2018; Yuan et al., 2019). An infected unit generates new secondary infections based on the probability density function of the generation time *g_w_* (***θ**_g_*). When a disease invades a new area, the index outbreak becomes the first generation. The index outbreak then generates *R*_1_ new cases (second generation) with illness onset spread out over time according to the distribution *g_w_*. The second generation cases *R*_1_ subsequently generate *R*_2_ new cases according to the same distribution *g_w_*. We derived the expected total number of generated cases as: *R*_1_(*g_w_* + *R*_2_(*g * g*)_*w*_) using a maximum of three generations and assuming independent and identical transmission events. The symbol “*” stands for the convolution function, as described by:

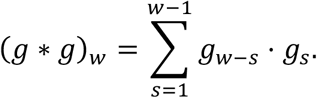

This method can be easily expanded to include a larger number of generations. We defined the force of infection in the following normalized form:

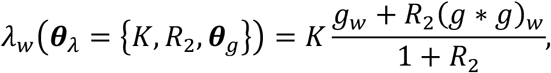

where we eliminated *R*_1_ from both the numerator and denominator as it was a common multiplier, and introduced a normalizing constant *K* that represents the expected total number of infected units.

We identified the lognormal distribution as the best candidate distribution for *g_w_* among gamma, lognormal, and Weibull distributions by comparing Akaike Information Criterion values for each model fit with the lowest value preferred. We then used Bayesian inference for the generation-dependent model with a lognormally distributed generation time to estimate the model parameters ***θ**_λ_* including their uncertainty levels.

We determined that a three-generation process captured the dynamics of the epidemic sufficiently well (Appendix Figure 1C), and the best-fitted lognormal distribution estimated the mean generation time at 7.54 weeks (95% CI: 6.60–8.90) with an SD of 1.73 weeks (95% CI: 0.92–3.06) (Appendix Figure 1D). Our model fit also indicated that the between-farm reproduction number was below one for the second generation *R*_2_, at which point the outbreak had been fully recognized by the authorities and control measures were implemented (median value: 0.43, 95% CI: 0–0.88).

### Appendix II. Underascertainment of cases results in overestimated mean transmission distance

We performed numerical simulations to quantify the effect of underascertainment on the estimated mean transmission distance we calculated using the spatiotemporal casedistribution method developed by Salje and colleagues (Salje et al., 2016). In order to do so, we used the Animal Disease Spread Model (ADSM)—a stochastic, spatial, state-transition simulation model for the spread of highly contagious diseases of animals (ADSM Development Team, 2019; Harvey et al., 2007). We used a sample population of pig farms provided by the developers for default setting: 461 units were distributed over a circular area with radius of 600 km. The model parameters were chosen to simulate a situation close to a real spread of ASF. We set the incubation period distribution to a Gamma distribution with a mean of 6.3 days and SD of 1.3 days (Olesen et al., 2017). The infectious period was assigned to a gamma distribution with a mean of 9.15 days and SD of 1.92 days, but we approximated the generation time distribution, defined as a convolution of the incubation period distribution and infectiousness period distribution, using a gamma distribution with resulting mean of 9.45 days (95% CI: 6.04–13.42) and SD of 2.30 days (95% CI: 1.23–3.50). The direct (within-pen) and indirect (between-pen) contact rates were set to 2.62 and 0.99, respectively (Hu et al., 2017). As ASFV is highly pathogenic, we set the probability of direct and indirect successful transmission to be 1.0. ADSM only considers airborne transmission of the virus, which we modelled using an exponential decay function with effective distance of 10 km. No control measures were considered in the simulations, which resulted in nearly 100% infection of the population. A sample run over 129 days resulted in infection of 440 out of 461 virtual swine farms (Appendix Figure 2AB).

From the results of the simulation, we estimated the mean transmission distance without any underascertainment to be 58 km, thereby confirming that the removal of geographic information for even large fraction of cases does not affect the obtained estimate (Salje et al., 2016)—see dashed orange in Appendix Figure 2C. However, we also found that the complete removal of outbreaks from the dataset results (reflecting underascertainment of incidence of infection) resulted in substantial overestimation of the mean transmission distance—see solid blue in Appendix Figure 2C. We thus argue that underascertainment of cases may significantly contribute to overestimation of the mean transmission distance when using the spatiotemporal case-distribution algorithm developed by Salje and colleagues.

**Appendix Figure 1.**
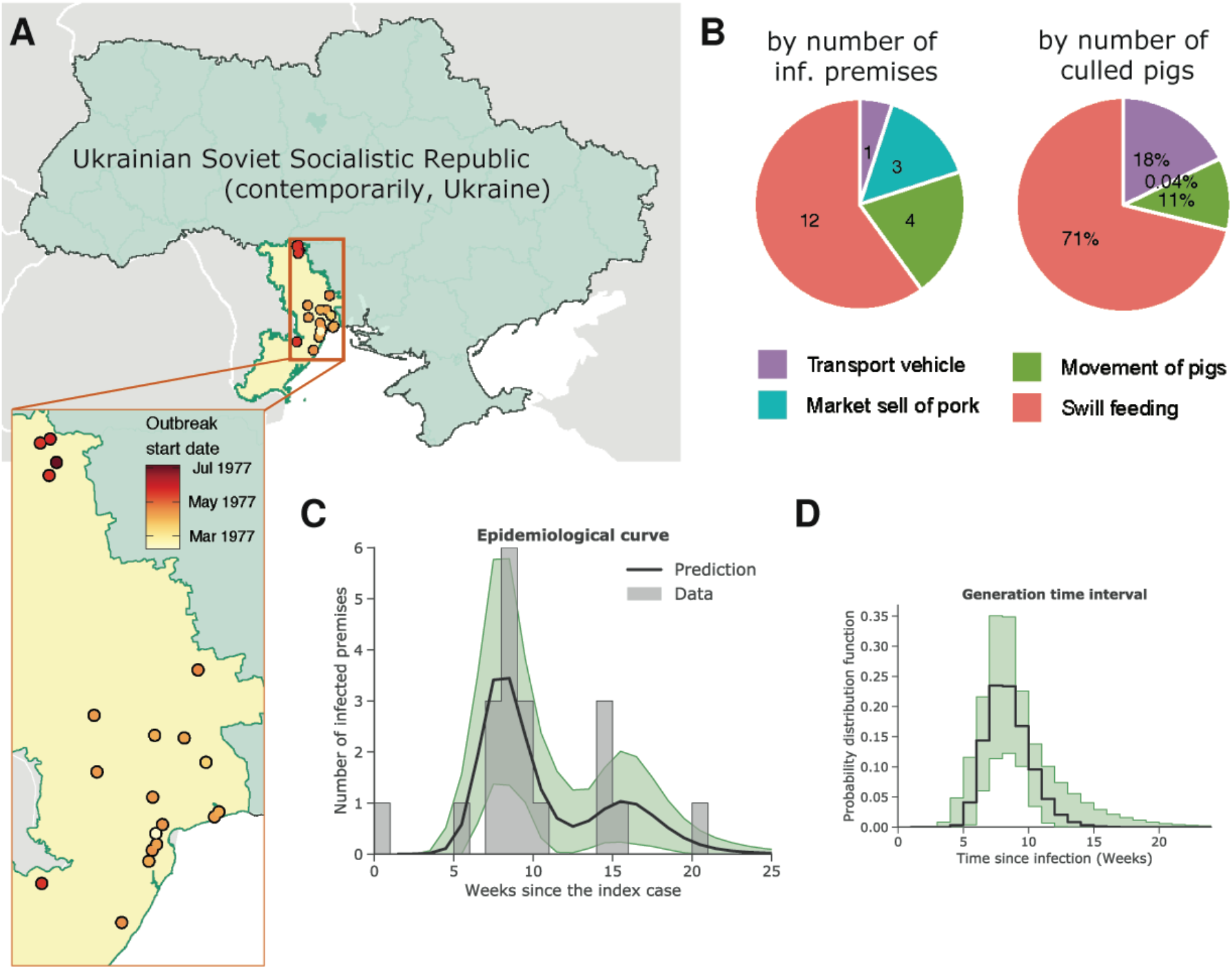
Analysis of a historical epidemic of ASF in Odessa region of Ukrainian Soviet Socialistic Republic (SSR) of the former Soviet Union, 1977, to determine the generation time distribution. (**A**) shows the map of Ukrainian SSR with highlighted locations of the reported outbreaks. (**B**) depicts the transmission routes determined as most likely by investigation of the epidemic. The left panel is respectively to the number of infected units, the right panel is respectively to the number of culled pigs: market sell of port products affected only backyard private farms. (**C**) shows the fitted multi-generational model to the data, whereas (**D**) the resulting generation time distribution. Shaded area in green indicates 95% credible interval.

**Appendix Figure 1—figure supplement 1.**
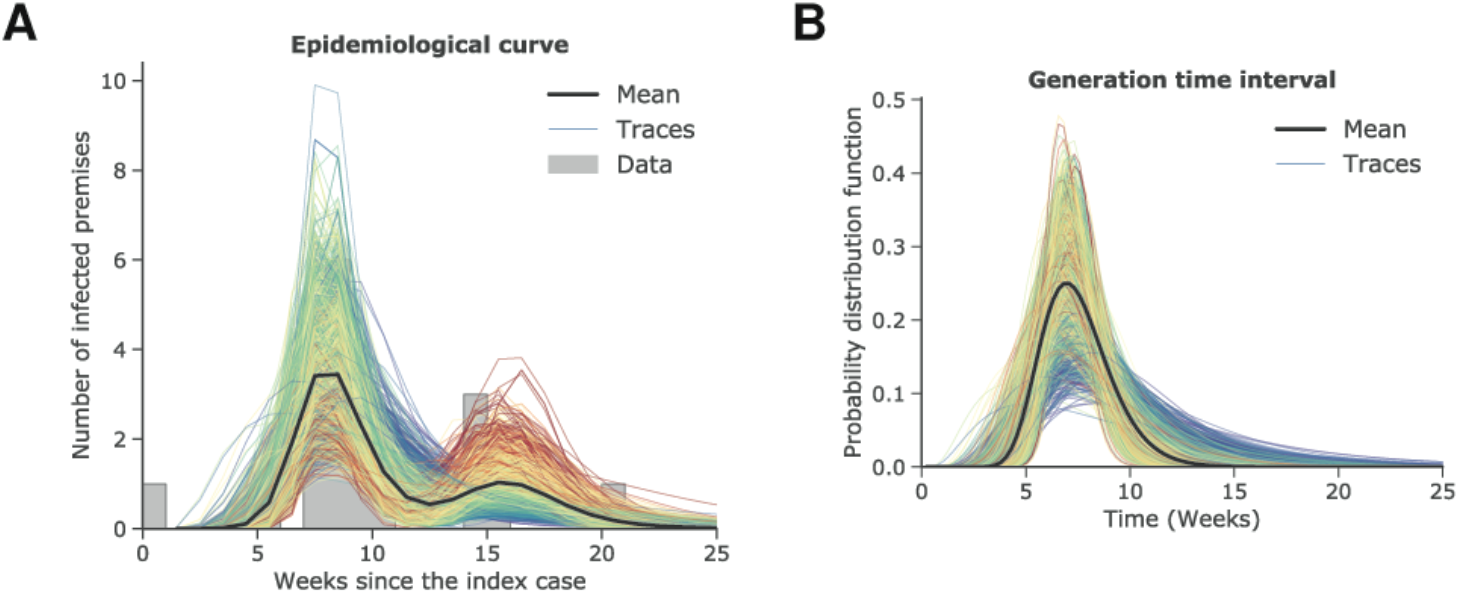
Trace plots of MCMC samples respective to the weekly number of cases (A) and generation time (B) show that both models with one and two generation of cases can be plausible. The colour code indicates the varying values of *R*_2_. When *R*_2_ is close to zero, the lines are blue, and the lines are red for high values of *R*_2_. Solid lines correspond to the mean values.

**Appendix Figure 2.**
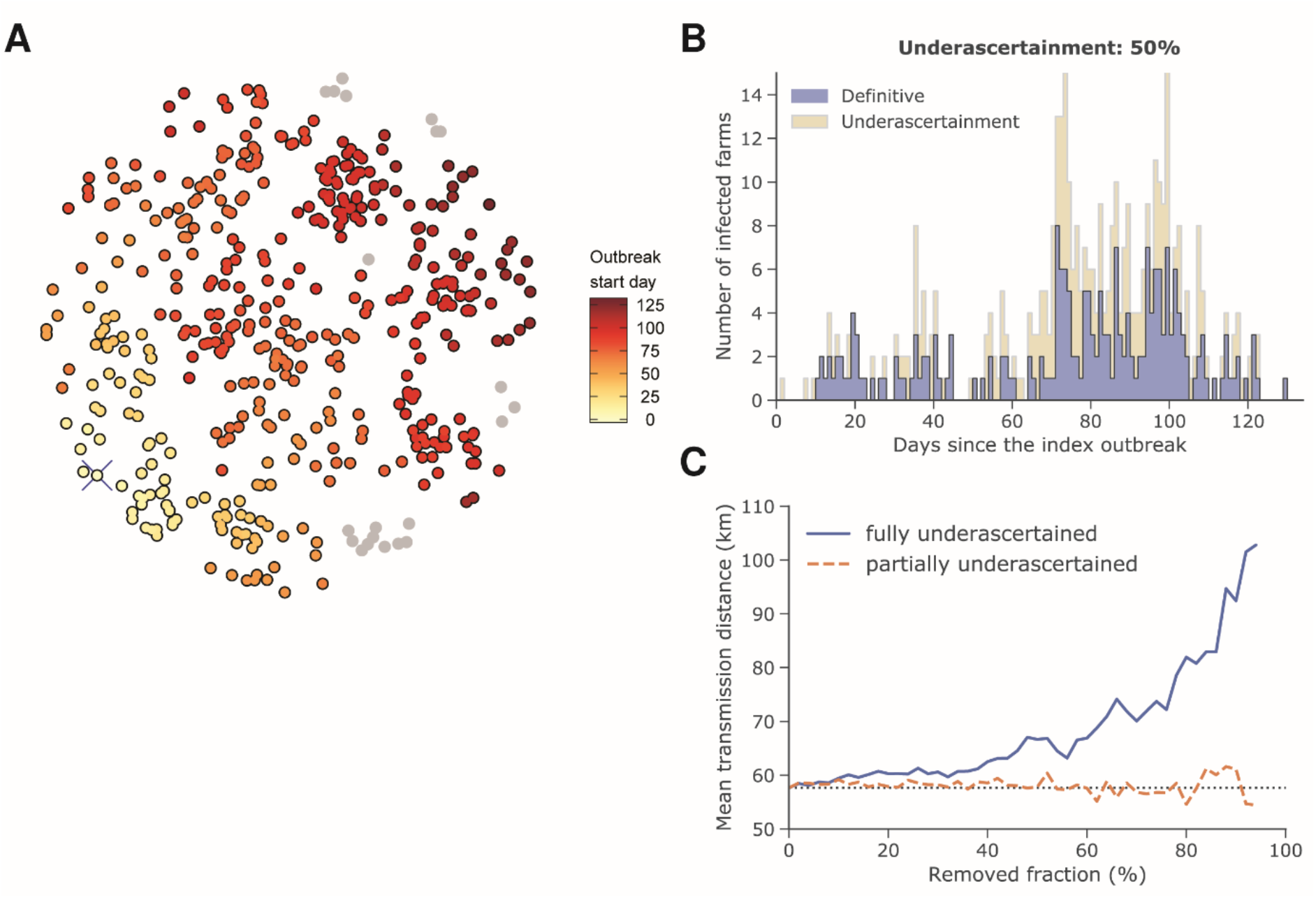
Simulated outbreak of ASF using Animal Disease Spread Model (ADSM Development Team, 2019). (A) shows the spatiotemporal spread of the disease. The crossed yellow dot in the bottom left-hand of the circle is the index case. Grey points represent uninfected farms. (B) depicts the epidemic curve. The dark bars represent definitive (reported) cases and the light bars represent partially or fully underascertained cases—i.e., cases missing spatial (geolocation) information or unreported cases that are missing both spatial and temporal information. (C) Estimation of the mean transmission distance for fully underascertained cases (solid line) or partially underascertained cases (dashed line). The dotted horizontal line is the estimate for the dataset with no underascertainment.

## Notes

### Competing Interest Statement

The authors have declared no competing interest.

